# Repurposing of mitochondria-targeted tamoxifen: Novel anti-cancer drug exhibits potent activity against major protozoan and fungal pathogens

**DOI:** 10.1101/2022.03.24.485593

**Authors:** Dominik Arbon, Kateřina Ženíšková, Karolína Šubrtová, Jan Mach, Jan Štursa, Marta Machado, Farnaz Zahedifard, Tereza Leštinová, Carolina Hierro-Yap, Jiri Neuzil, Petr Volf, Markus Ganter, Martin Zoltner, Alena Zíková, Lukáš Werner, Robert Sutak

## Abstract

Many of the currently available anti-parasitic and anti-fungal frontline drugs have severe limitations, including adverse side effects, complex administration, and increasing occurrence of resistance. The discovery and development of new therapeutic agents is a costly and lengthy process. Therefore, repurposing drugs with already established clinical application offers an attractive, fast-track approach for novel treatment options. In this study, we show that the anti-cancer drug MitoTam, a mitochondria-targeted analog of tamoxifen, efficiently eliminates a wide range of evolutionarily distinct pathogens *in vitro*, including pathogenic fungi, *Plasmodium falciparum*, and several species of trypanosomatid parasites, causative agents of debilitating neglected tropical diseases. MitoTam treatment was also effective *in vivo* and significantly reduced parasitemia of two medically important parasites, *Leishmania mexicana* and *Trypanosoma brucei*, in their respective animal infection models. Functional analysis in the bloodstream form of *T. brucei* showed that MitoTam rapidly altered mitochondrial functions, particularly affecting cellular respiration, lowering ATP levels, and dissipating mitochondrial membrane potential. Our data suggest that the mode of action of MitoTam involves disruption of the inner mitochondrial membrane, leading to rapid organelle depolarization and cell death. Altogether, MitoTam is an excellent candidate drug against several important pathogens, for which there are no efficient therapies and for which drug development is not a priority.

**Author Summary:** MitoTam, a mitochondrially targeted analog of tamoxifen, is a promising anti-cancer candidate drug acting by accumulating in and destabilizing cell mitochondria. In this study, we analyze its effect on a wide range of evolutionarily distinct and medically important pathogens. These include a) pathogenic fungi, *Candida albicans,* and *Cryptococcus neoformans,* ubiquitous opportunistic pathogens that cause life-threatening diseases in immunocompromised or immunologically deficient individuals; b) *Plasmodium falciparum*, the causative agent of human malaria; and c) several species of trypanosomatid parasites such as *Trypanosoma cruzi,* responsible for deadly Chagas disease in South America, *Trypanosoma brucei,* the causative agent of sleeping sickness in Africa, and *Leishmania, the* etiological agent of leishmaniasis, a spectrum of diseases ranging from usually self-healing but potentially disfiguring cutaneous and mucocutaneous leishmaniasis, to visceral leishmaniasis, which is invariably fatal if left untreated. We show that MitoTam efficiently kills these parasites in laboratory conditions and in the case of trypanosomes and leishmaniases suppress or at least slow down the infection in mouse model.

## Introduction

The ongoing search for novel treatment options to combat medically relevant parasitic protists (e.g. *Plasmodium*, *Trypanosoma* and *Leishmania spp.*) is driven by the need for more efficient, less toxic, and/or less expensive medications as well as by the emergence and spread of drug resistance [1–3]. Drug discovery and development have been facilitated in recent decades by advances in the fields of genetics, molecular biology, medicinal chemistry and the introduction of high throughput target-based, phenotypic and virtual screening strategies. Nevertheless, repurposing of drugs originally approved for other indications presents a strategy of particular interest for the implementation of novel treatments for neglected diseases [3]. As repurposed drugs are typically at least partly characterized or even approved for clinical use, both the time and cost of the drug development process are dramatically reduced. This is particularly appealing for drugs against neglected diseases with little financial incentive for commercial ‘*de novo’* drug discovery approaches.

Phosphonium salts are lipophilic cations with the ability to readily pass across phospholipid bilayers, and due to their delocalized positive charge, they accumulate at the interface of the inner mitochondrial membrane (IMM) and matrix according to the mitochondrial membrane potential (ΔΨ_m_). The extent of accumulation of lipophilic cations at the IMM follows the Nernst equation, according to which there is a tenfold increase in the concentration of lipophilic cations at the IMM- matrix interface for every ∼60 mV increase in ΔΨ_m_ [4]. Phosphonium vectors have been employed for efficient and selective mitochondrial delivery of various cargo molecules such as therapeutic antioxidants [5, 6], liposomes [7], functional probes [8, 9], anti-microbial, anti-fungal and anti-parasitic drugs [10–16] and anti-cancer treatments [10,17,18].

MitoTam, mitochondrially targeted cationic triphenylphosphonium vector (TPP^+^) conjugated with tamoxifen, is a promising anti-cancer candidate drug acting by mitochondrial destabilization [19]. It was originally developed to selectively accumulate the pharmacophore tamoxifen in the mitochondria proportionally to ΔΨ_m_. The compound is well tolerated in the mouse model and recently underwent a phase 1/1b clinical trial in human advanced-stage cancer patients with favorable outcome (MitoTam-01 trial; EudraCT 2017-004441-25). Its molecule consists of three parts: i) the functional pharmacophore residue, tamoxifen, a drug that has been used for decades to treat early and advanced hormone-dependent breast cancer [20], ii) the TPP^+^ for mitochondrial targeting, and iii) the ten-carbon linear alkyl linker tethering tamoxifen to the TPP^+^ moiety. The compound efficiently kills breast cancer cells and suppresses tumors progression, including treatment-resistant carcinomas with high Her2 protein levels (Her2^high^), for which the original precursor compound tamoxifen was ineffective. Importantly, MitoTam exhibits low toxicity towards non-malignant cells [19]. Similar to the mitochondria-mediated anti-cancer effects of tamoxifen, MitoTam modulates or alters multiple mitochondrial processes including the function of the respiratory complex I (CI; NADH:ubiquinone dehydrogenase) [19,21–23]. The superior efficacy of MitoTam compared to tamoxifen is due to its accumulation at the interface of the mitochondrial matrix and IMM that leads to suppression of CI-dependent respiration, disruption of respiratory supercomplexes, rapid dissipation of the ΔΨ_m_, increased production of reactive oxygen species (ROS), and ultimately cell death [19]. MitoTam is also effective in the treatment of the syngenic renal cancer murine model, especially in combination with immunotherapy [23]. In addition, MitoTam selectively eliminates senescent cells and is, therefore, a candidate for the treatment of age-related disorders [22, 24].

In this work, we tested the potential activity of MitoTam against a wide range of parasitic protists and pathogenic fungi (Table S1). These models were selected for their medicinal and economic relevance and for their suitability as cellular models for research of mitochondrial function. We chose parasites from the Kinetoplastida group: *Trypanosoma brucei* as a causative agent of human and animal African trypanosomiases and a highly tractable model organism; *Trypanosoma cruzi*, an intracellular parasite responsible for Chagas disease; and *Leishmania spp.*, the etiological agent of a spectrum of disorders ranging from self-healing cutaneous lesions to potentially fatal visceral diseases [25]. We also tested the effect of MitoTam on *Plasmodium falciparum*, a parasite responsible for most malaria-related deaths in humans [26]; on the pathogenic fungi *Candida albicans* and *Cryptococcus neoformans*, widespread opportunistic pathogens causing life-threatening diseases in immunocompromised individuals [27, 28]; on the amphizoic amoebae *Naegleria fowleri* and *Acanthamoeba* spp., whose infections lead to rare diseases with extremely high mortality rate [29, 30]; and on *Giardia intestinalis* and *Trichomonas vaginalis*, anaerobic parasites with reduced mitochondria, causative agents for widespread intestinal and urogenital infections [31–33]. Here, we show that the anti-cancer drug candidate MitoTam efficiently and selectively inhibits viability of a number of these pathogens *in vitro*. In pilot experiments using mouse models of infection with *T. brucei* and *L. mexicana*, MitoTam reduced parasite burdens and significantly prolonged survival for *T. brucei*-infected animals. Furthermore, we demonstrate in *T. brucei* that the trypanocidal activity of MitoTam relies, at least partly, on rapid disruption of IMM. Together, our data provide strong evidence that MitoTam is an excellent candidate for further development as an anti-infective, especially for targeting several neglected diseases.

**Table 1:** MitoTam inhibits the growth of several pathogenic microorganisms in vitro. Mean EC50 values for MitoTam, tamoxifen (the functional pharmacophore of MitoTam) and specific anti- microbial drugs used as positive controls for each tested organism derived from dose-response curves are given in µM (n>3, ± s.d.).

| Pathogen | MitoTam |  | Tamoxifen |  | Control compound |  |  |
| --- | --- | --- | --- | --- | --- | --- | --- |
|  | EC<sub>50</sub> [μM] | SD | EC<sub>50</sub> [μM] | SD |  | EC<sub>50</sub> [μM] | SD |
| *Trypanosoma brucei* BSF | 0,02 | ±0.00 | 7,20 | ±1.47 | amphotericin B | 1,09 | ±0.09 |
| *Trypanosoma brucei* PCF | 0,16 | ±0.03 | ≥ 10 |  | amphotericin B | 0,24 | ±0.06 |
| *T. brucei gambiense* BSF | 0,03 | ±0.00 | ≥ 10 |  | pentamidine | 0,00 | ±0.00 |
| *Trypanosoma cruzi* (epimastigotes) | 1,55 | ±0.04 | ≥ 10 |  | benznidazole | ≥ 10 |  |
| *Leishmania mexicana* (amastigotes) | 0,35 | ±0.05 | 7,93 | ±0.59 | amphotericin B | 0,33 | ±0.08 |
| *Leishmania major* (promastigotes) | 0,60 | ±0.10 | ≥ 10 |  | amphotericin B | 0,06 | ±0.02 |
| *Plasmodium falciparum* (erythrocytic stage) | 0,03 | ±0.01 | 6,37 | ±0.11 | chloroquine | 0,01 | ±0.00 |
| *Candida albicans* | 0,56 | ±0.07 | ≥ 10 |  | amphotericin B | 0,37 | ±0.01 |
| *Cryptococcus neoformans* | 0,61 | ±0.12 | 7,28 | ±1.97 | amphotericin B | 0,12 | ±0.05 |
| *Naegleria fowleri* | ≥ 10 |  | ≥ 10 |  | amphotericin B | 0,08 | ±0.01 |
| *Acanthamoeba* spp. | 1,95 | ±0.23 | 9,93 | ±0.58 | amphotericin B | 6,17 | ±0.70 |
| *Giardia intestinalis* | ≥ 10 |  | ≥ 10 |  | benznidazole | 6,22 | ±0.66 |
| *Trichomonas vaginalis* | ≥ 10 |  | ≥ 10 |  | metronidazole | 6,54 | ±0.45 |

## Results

### Low levels of MitoTam are lethal to a variety of pathogenic eukaryotic microorganisms

The mitochondrion is a promising drug target because of its pivotal role in cellular metabolism, proliferation, and cell death signaling [34–36]. We therefore investigated the effect of MitoTam, a mitochondria-targeted tamoxifen derivative, on a wide range of parasitic protists and yeasts. MitoTam proved to have a significant growth inhibitory effect for the majority of these pathogens (Table 1) exhibiting nanomolar potencies against *P. falciparum*, *T. b. brucei* and *Leishmania spp.* Notably, the half-maximal effective concentration (EC_50_) values are considerably lower than corresponding cytotoxicity values reported for various mammalian cells (ranging from 0.65-4.5 µM) [19], indicating high selectivity. All pathogens were also challenged with the parental compound tamoxifen lacking the TPP^+^ vector for mitochondrial targeting, with efficacy significantly lower compared to MitoTam (Table 1). This is consistent with similar comparisons reported for the anti-cancer activity of the two molecules [19]. Each dose-response analysis was validated using a known, active compound against individual pathogens tested, providing values consistent with published data (Table 1).

The highest efficacy of MitoTam was detected for the bloodstream form (BSF) of *T. brucei*. This value was approximately 30-fold greater compared to Her2^high^ cancer cells (0.02 µM *vs* 0.65 µM) [19]. Interestingly, MitoTam showed low nanomolar efficacy (0.03 µM) against the erythrocytic stage of *P. falciparum*, which is protected inside the parasitophorous vacuole within the host erythrocyte [37], suggesting that MitoTam is able to cross several membranes before reaching its target. Furthermore, MitoTam was found to inhibit the growth of the *T. brucei* procyclic form (PCF), axenically grown *Leishmania mexicana* amastigotes and *L. major* promastigotes, and the two pathogenic fungi *Candida albicans* and *Cryptococcus neoformans* at sub-micromolar EC_50_ values. *T. cruzi* epimastigotes also responded to MitoTam treatment with low-micromolar EC_50_, as did *Acanthamoeba spp.*, while no effect was observed on *Naegleria fowleri* proliferation. Also, *Trichomonas vaginalis* and *Giardia intestinalis*, two anaerobic parasites that possess mitochondrion-related organelles characterized by the absence of membrane-bound electron transport chain and therefore by the lack of mitochondrial respiration [33], were insensitive to MitoTam treatment. Overall, MitoTam effectively eliminated a range of clinically important, evolutionarily distinct pathogens including intracellular parasites.

### MitoTam efficiently eliminates *L. major* and *L. infantum* intracellular amastigotes

The high efficacy of MitoTam against *L. mexicana* axenic amastigotes (EC_50_ 0.35 ± 0.05 μM) prompted us to investigate the ability of MitoTam to eliminate the intracellular form of the parasite in a macrophage infection model. Murine macrophage cells (J774A.1) were infected with *L. major* or *L. infantum* promastigotes and exposed to different concentrations of MitoTam (0-25µM). After controlled lysis of the infected macrophages, the released parasites were quantified using a resazurin-based viability assay [38]. In this intramacrophage assay, treatment with MitoTam eliminated intracellular amastigotes of *L. major* and *L. infantum* with EC_50_ values of 175 ± 51 nM and 293 ± 29 nM, respectively (Fig. S1). This result is highly encouraging since the parasites reside in a parasitophorous vacuole, a phagolysosome-like structure with low pH and three membranes that make it difficult to effectively target the intracellular *Leishmania* amastigotes *in vivo* [38].

### MitoTam treatment alleviates parasitemia leading to prolonged survival of mice infected with *T. brucei* and to reduced frequency and lesion size caused by *L. mexicana* infection

Activity for a compound series detected in the intramacrophage assay usually translates into activity in an animal infection model [39, 40], apart from potential issues with pharmacokinetics. Hence, we next tested the ability of MitoTam, which is well tolerated in BALB/c mice [19], a mouse infection model of both *L. mexicana* and *T. brucei.* MitoTam dosing regimen was based on the published data [19, 22].

For the *T. brucei* infection model, survival analysis revealed that MitoTam intravenous (IV) administration at doses of 3 mg/kg body weight (bw) on days 3 and 5 post-infection without further treatment delayed the death of *T. brucei* infected animals, which succumb to the infection by eight days (Fig. 1A).

**Figure 1:**
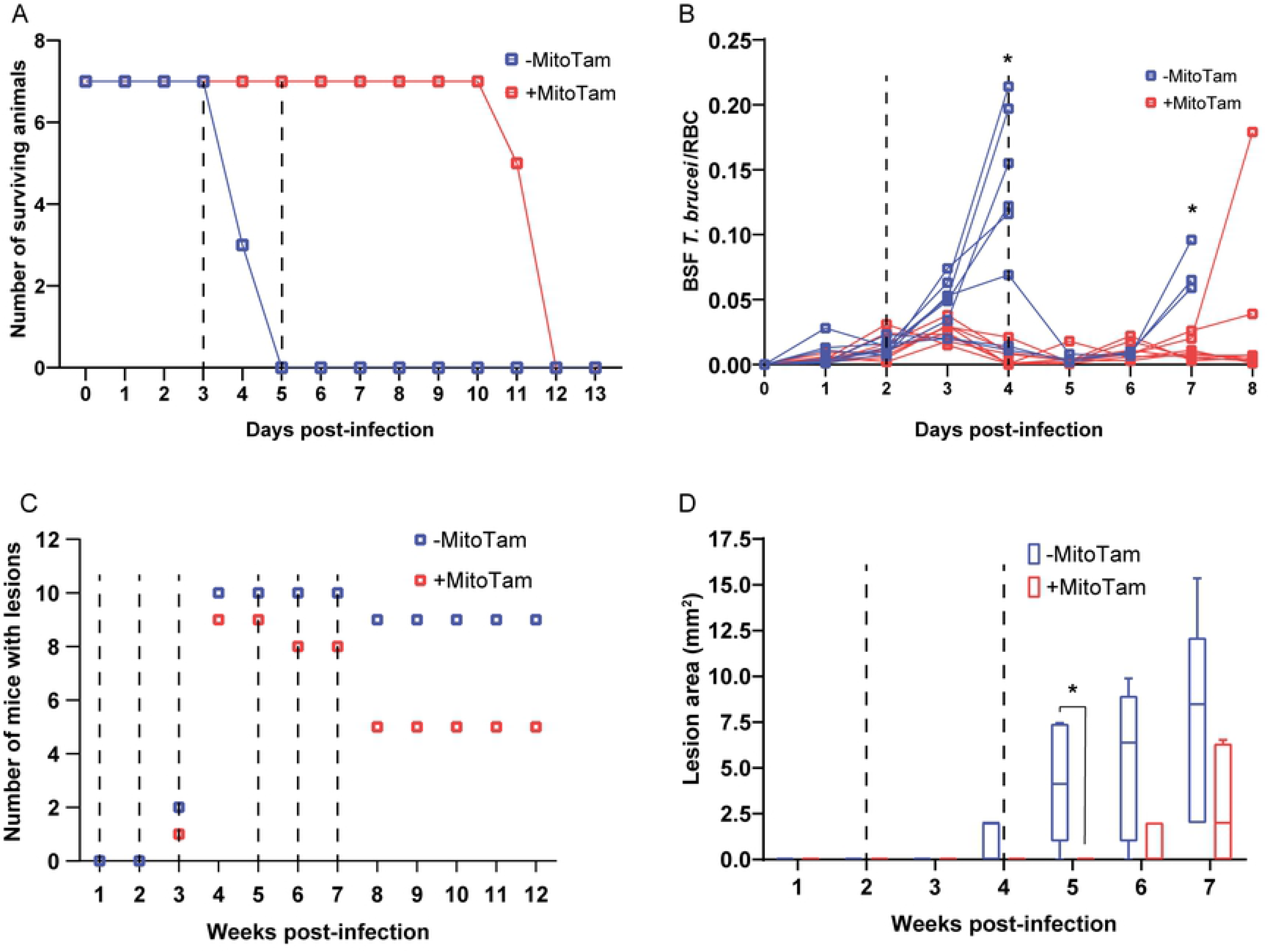
MitoTam is effective against *T. brucei* and *L. mexicana* parasites *in* mouse infection models. **A)** Survival rate of BALB/c mice infected with *T. brucei* was monitored daily for 13 days. Half of the infected mice were injected IV with MitoTam (+MitoTam) on days 3 and 5 (dashed lines) post-infection (3 mg/kg bw). The number of surviving mice from MitoTam treated group (red line) as well as from untreated control group (-MitoTam) (blue line) is plotted against the time (in days). **B)** Parasitemia in BALB/c mice infected with *T. brucei* was evaluated microscopically from blood smears. On day 2 and 4 post-infection (dashed lines), eight mice were IV injected with MitoTam (3 mg/kg) (+MitoTam). The ratio of *T. brucei* cells to red blood cells (RBC) was calculated daily and used to quantify the parasitemia levels of each animal from the MitoTam treated group (red line) and the respective untreated control group (-MitoTam) (blue line). Statistically significant differences between the two groups are indicated with an asterisk (unpaired t-test, * p < 0.05). **C)** Formation of dermal lesions in twenty BALB/c mice was followed for 12 weeks after infection with *L. mexicana*. As indicated (dashed lines), half of the infected animals were injected IP with MitoTam (+MitoTam) at 1, 2, 3, 5, 6 and 7 weeks post-infection. The appearance of lesions was monitored (+MitoTam, red) and compared to the number of lesions of animals from the untreated control group (-MitoTam, blue). **D)** Size of dermal lesions was recorded in ten BALB/c mice infected with *L. mexicana*. Five of the infected animals were treated with MitoTam (+MitoTam) at weeks 2 and 4 post-infection as indicated (dashed lines). Throughout the course of the infection, the lesion areas of each animal were measured weekly. The values were averaged for the MitoTam treated group (+MitoTam, in red) and untreated control group (-MitoTam, in blue). Mean values and error bars representing the standard deviations are shown, statistically significant differences are indicated with an asterisk (paired t-test, *p < 0.05).

In a second experiment, we followed *T. brucei* parasitemia levels in infected animals (Fig. 1B). Automated analysis of blood smears revealed the presence of *T. brucei* parasites in samples from mice untreated and treated on days 2 and 4 post-infection, showing that MitoTam-treated animals exhibited a significantly lower parasite load (Fig. 1B). This explains the longer survival of treated animals compared to the untreated controls (Fig. S2).

The effect of MitoTam administration (3 mg/kg bw, intraperitoneally (IP)) in the *L. mexicana* infection model was assessed by monitoring frequency and size of dermal lesions. As shown in Fig. 1C, lesions occurred less frequently in MitoTam-treated BALB/c mice than in the untreated controls during the 12-week course of the experiment (50% vs. 90% of mice at weeks 8-12 post-infection). In a separate experiment, MitoTam at two doses reduced the size of lesions caused by *L. mexicana* in the weeks following infection (Fig. 1D).

Collectively, these data demonstrate that MitoTam is highly effective against *T. brucei* and *L. mexicana* parasites *in vivo*. Notably, we did not observe any notable adverse effects of MitoTam on the treated animals.

### MitoTam treatment causes alteration of *T. brucei* mitochondrial proteome and leads to upregulation of multidrug efflux pumps in *C. albicans*

The potent anti-parasitic properties *in vitro* and the promising *in vivo* experiments prompted us to explore the MitoTam mode of action. First, we performed comparative whole-cell proteomic analysis of control and MitoTam-treated cells for two parasitic model organisms, *C. albicans* and *T. brucei.* In order to restrict indirect, secondary impact resulting from cell death, we chose short exposure times: yeast cells were treated with 4.4 μM MitoTam (⁓7x EC_50_) for 12 h, while *T. brucei* BSF cells were incubated with 100 nM MitoTam (5x EC_50_) for 14 h. Proteomic data were processed by label-free quantification in MaxQuant [41] and statistically analyzed in Perseus [42, 43]. In *C. albicans*, out of 1,950 detected protein groups, the abundance of only 1.69% of the quantified proteins was significantly altered (Table S2) (Fig. 2A), with two homologs of multidrug efflux pumps being among the most upregulated genes (≥26 times).

**Figure 2:**
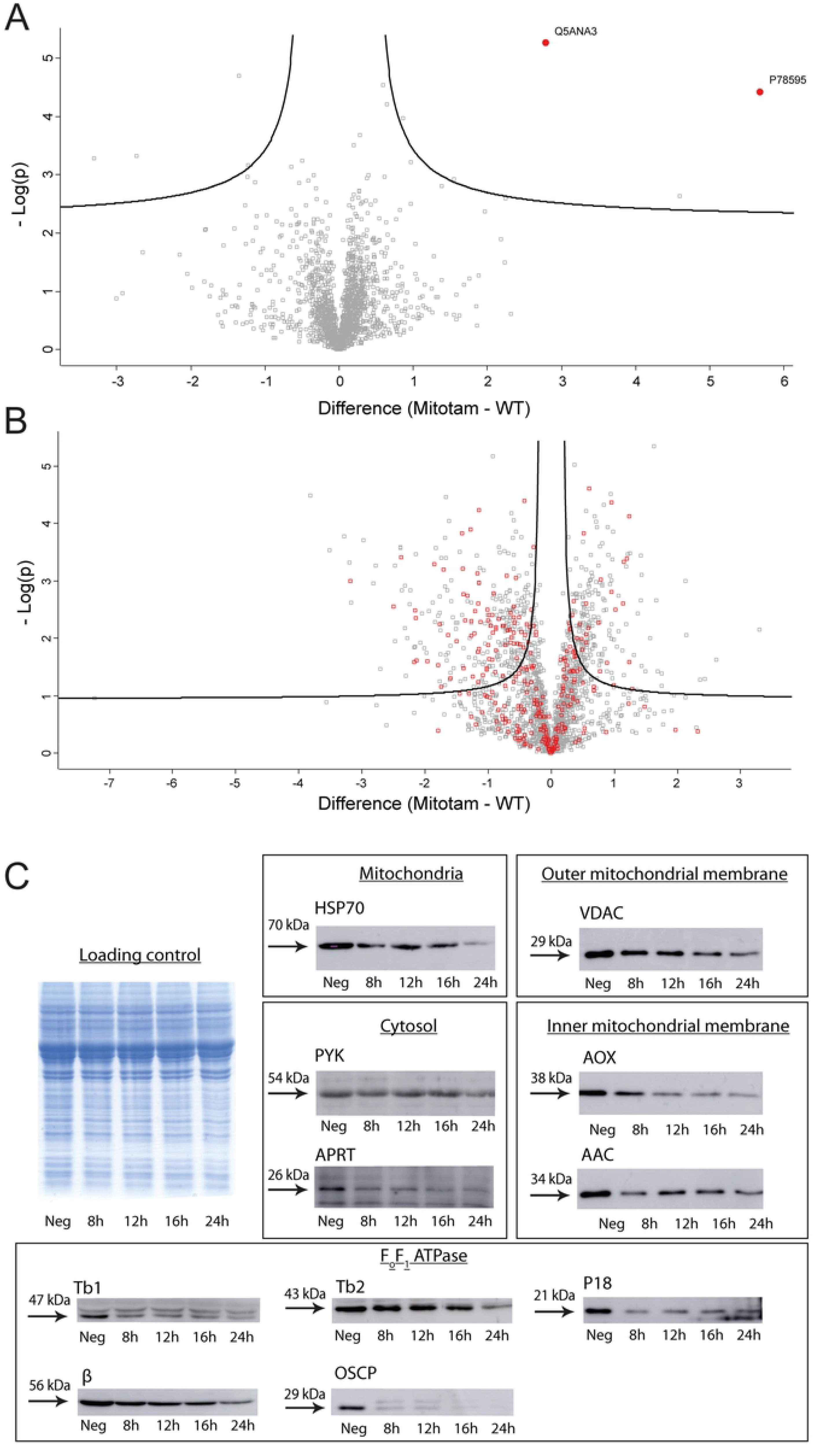
Global proteome changes upon MitoTam exposure. Shown are volcano plots of the *t*-test difference from label-free quantification from triplicate experiments, plotted against the respective −log_10_-transformed *p* values. Untreated control cells were compared to cells treated with MitoTam for *C. albicans* (A) or BSF *T. brucei* (B). **A)** Highlighted points are *C. albicans* significantly upregulated efflux pumps (red dots): P78595 - Multidrug resistance protein CDR2; Q5ANA3 - Pleiotropic ABC efflux transporter of multiple drugs CDR1. **B)** *T. brucei* mitochondrial proteins are in red. **C)** Western blot analyses of whole-cell lysates harvested from BSF *T. brucei* cells treated with 100 nM MitoTam for 0-24 hours as indicated. Equal amounts of total protein were loaded in each lane. A Coomassie stained gel was used as a control for equal loading. The immunoblots were probed with antibodies against cytosolic pyruvate kinase (PYK) and adenine phosphoribosyltransferase (APRT) and mitochondrial heat shock protein 70 (HSP70), proteins associated with the outer mitochondrial membrane (VDAC), inner mitochondrial membrane (AOX, AAC), and subunits of the F_o_F_1_-ATP synthase (β, p18, OSCP, Tb1, Tb2). The relevant masses of the protein molecular weight are indicated.

In contrast, analysis of the *T. brucei* proteome revealed substantial changes after treatment with MitoTam, with 26.3% of all identified proteins (2,063) significantly altered (Table S3) (Fig. 2B). Data analyses did not reveal any specific single pathway or protein as a direct target of MitoTam. Since MitoTam specifically affects mitochondrial functions in various cells [19,22,23], we analyzed enrichment of mitochondrial proteins using a *T. brucei* mitoproteome database [44]. While mitochondrial proteins accounted for 14.3% of all detected proteins, we observed an almost 2-fold enrichment (27.6%) in the significantly altered subset of proteins, with 20.8% and 32.1% in the increased and decreased cohort, respectively. These results indicate that MitoTam treatment induces preferentially a decrease in the levels of various mitochondrial proteins. To validate these results, we analyzed *T. brucei* whole cell lysates from cultures harvested at 4 different time points of MitoTam exposure (8, 12, 16, and 24 h) by immunoblotting with antibodies against 8 mitochondrial marker proteins and cytosolic pyruvate kinase as a control (Fig. 2C). Consistent with the *T. brucei* proteomics analysis, we detected significant abundance decrease for seven mitochondrial proteins after 8 h of MitoTam treatment.

### MitoTam treatment leads to disruption of the mitochondrial inner membrane and rapid loss of ΔΨ_m_ **in the bloodstream form of *T. brucei*.**

MitoTam was reported to directly impact mitochondrial function in renal and breast cancer cells, tumors, and senescent cells [19,22,23]. In line with this, our proteomic analysis indicated that MitoTam treatment perturbs the mitochondrial proteome in *T. brucei*. *T. brucei* BSF cells lack cytochrome-mediated mitochondrial electron transport chain, respiring exclusively via an alternative pathway consisting of mitochondrial glycerol-3-phosphate dehydrogenase and trypanosome alternative oxidase (AOX), and sustaining the ΔΨ_m_ by reversed activity of F_o_F_1_-ATP synthase [45–47]. Importantly, the cancer molecular target of MitoTam, respiratory CI [19] was shown to be dispensable for BSF cells [48], yet our data show that the drug efficiently eliminates these parasites.

To gain deeper insight into the anti-parasitic mode of action of MitoTam, we investigated its effect on mitochondria in this model protist. When comparing the growth of untreated cells with cells treated with two different concentrations of MitoTam, we found that growth of *T. brucei* cells was significantly inhibited after 24 h (40 nM, 2x EC_50_) or 12 h (100 nM, 5x EC_50_) (Fig. 3A). To establish a timeline for further experiments, we performed a live/dead cell assay using the cell-impermeant dye Sytox. This result shows that despite the reduced growth rate, 82.4 % and 71% cells were still viable upon 40 nM and 100 nM MitoTam treatment, respectively, after exposure to the drug for 16 h. At 24 hours of treatment, the viability was further decreased to 67.3% and 63.6% at 40 and 100 nM MitoTam, respectively (Fig. 3B)). This was accompanied by increased level of cellular ROS of cells treated with 100 nM MitoTam for 16 h (Fig. 4A).

**Figure 3:**
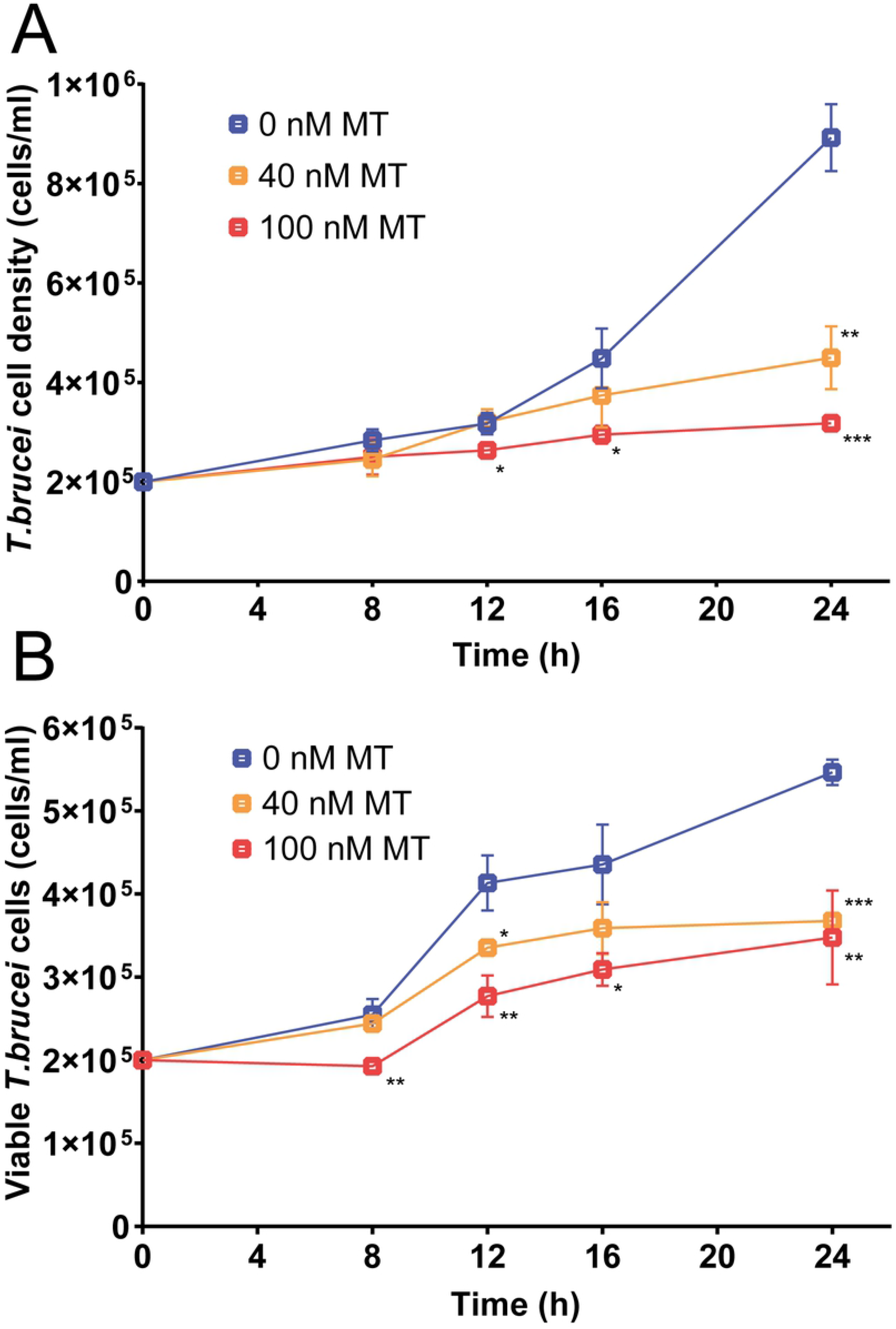
MitoTam affects *T. brucei* viability and alters the mitochondrial function of bloodstream *T. brucei* cells. **A)** Growth curve of BSF *T. brucei* cultures was evaluated in unstained cells by flow cytometry. Untreated control BSF cells (0 nM MT, blue) were compared with cells incubated with 40 nM MitoTam (40 nM MT, orange) and 100 nM MitoTam (100 nM MT, red). Statistically significant differences are indicated with an asterisk (paired t-test, *p < 0.05, **p < 0.01, ***p < 0.001) (mean ± s.d., n=3). **B)** Staining with the cell impermeant nucleic acid dye Sytox was used to distinguish viable (Sytox negative) and dead (Sytox positive) BSF *T. brucei* cells. Concentrations of Sytox negative, live cells are plotted. Untreated control cultures (0 nM MT, blue) were compared with cultures incubated in the presence of 40nM MitoTam (40 nM MT, orange) and 100 nM MitoTam (100 nM MT, red) (paired t-test, *p < 0.05, **p < 0.01, ***p < 0.001) (mean ± s.d., n=3).

**Figure 4:**
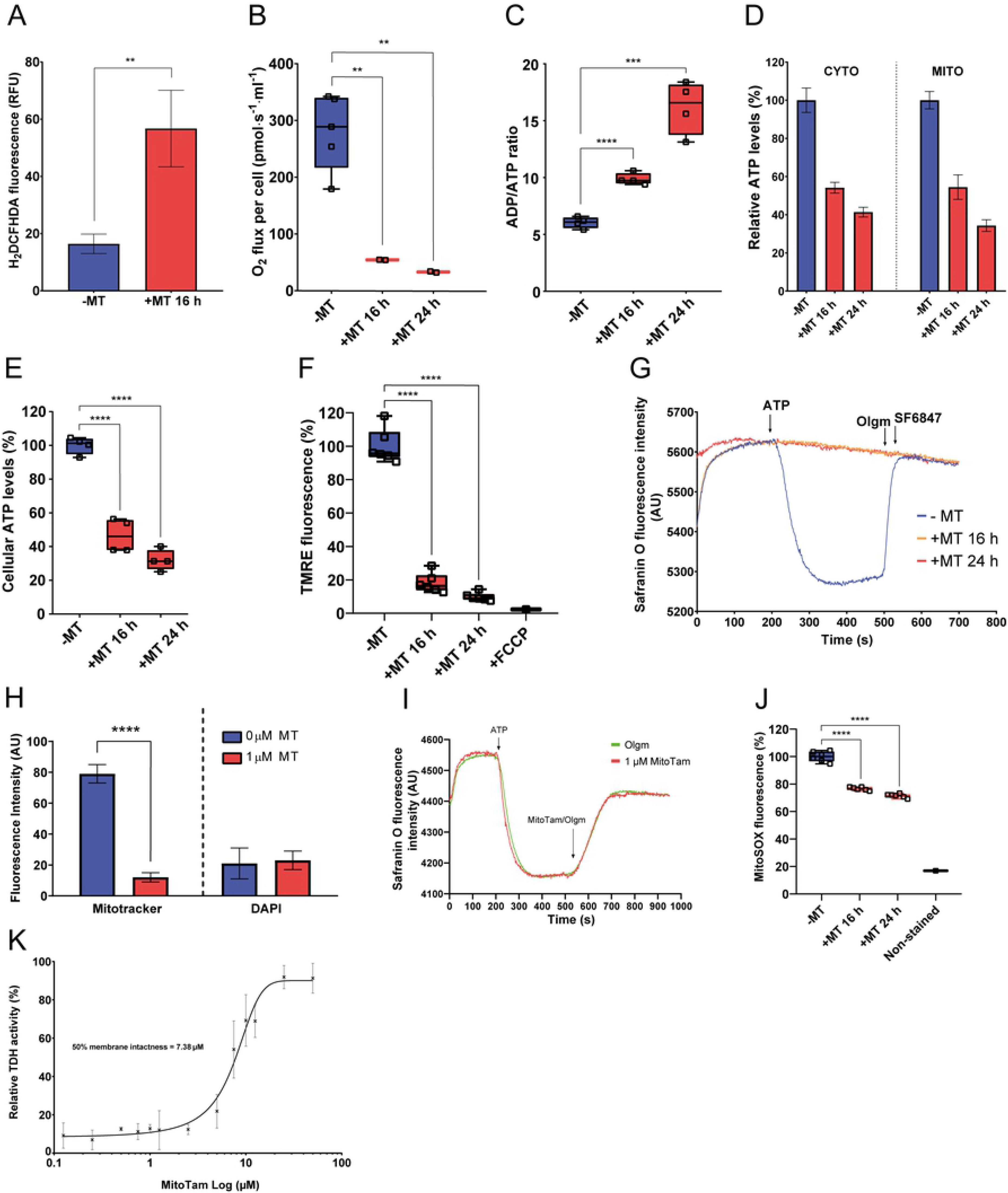
Overall effect of MitoTam on the *T. brucei* cell. **A)** Production of intracellular ROS in BSF *T. brucei* was quantified by flow cytometry using DCFH-DA detection reagent in untreated control cells (-MT, blue) and cells treated with 100 nM MitoTam for 16 hours (+MT 16 h, red) (mean ± s.d., *n* = 3, **p < 0.01). **B)** The O_2_ flux per cell using high-resolution respirometry after addition of glycerol-3-phosphate was determined in BSF *T. brucei* untreated control cells (-MT, blue) and BSF cells treated with 100nM MitoTam for 16 hours (+MT 16 h, red) and 24 hours (+MT 24 h, red) (mean ± s.d., *n* = 3, **p < 0.01). **C)** Relative ADP/ATP ratio analyzed using a bioluminescence assay kit in BSF *T. brucei* untreated control cells (-MT, blue) and BSF cells treated with 100 nM MitoTam for 16 hours (+MT 16 h, red) and 24 hours (+MT 24 h, red) (mean ± s.d., *n* = 3, ***p < 0.001, ****p < 0.0001). **D)** Cytosolic and mitochondrial ATP levels were assessed in transgenic BSF *T. brucei* cell lines expressing firefly luciferase. Untreated control cells (-MT, blue) were compared with cells treated with 100 nM MitoTam for 16 hours (+MT 16 h, red) and 24 hours (+MT 24 h, red). Data were normalized to the values of the untreated control cells and expressed as a percentage (mean ± s.d., *n* = 4). **E)** Total cellular ATP levels in BSF *T. brucei* were determined using a bioluminescence assay kit. Data from cultures treated with MitoTam for 16 hours (+MT 16 h, red) and 24 hours (+MT 24 h, red) were normalized to the respective values of the untreated control cells (-MT, blue) and expressed in percentage (mean ± s.d., n = 4), ****p <0.0001). **F)** Flow cytometry of TMRE stained cells was used to determine ΔΨ_m_ of BSF *T. brucei*. Data from cultures treated with 100 nM MitoTam for 16 hours (16 h +MT, red) and 24 hours (+MT 24 h, red) were normalized to the respective values of the untreated control cells (-MT, blue) and expressed in percentage. Uncoupler FCCP was added as a control for ΔΨ_m_ depolarization (mean ± s.d., n = 6, ****p<0.0001). **G)** *In situ* ΔΨ_m_ was measured in digitonin-permeabilized BSF *T. brucei* cells stained with Safranin O dye. Where indicated, the F_o_F_1_-ATP synthase substrate – ATP, the F_o_F_1_-ATP synthase inhibitor - oligomycin (Olgm) and the protonophore SF6847 were added. Representative trace from the measurement of untreated control cells (-MT, blue) in comparison with cells treated with 100nM MitoTam for 16 hours (+MT 16 h, orange) and 24 hours (+MT 24 h, red) is shown. **H)** Automated microscopic analysis of BSF *T. brucei* cells stained with Mitotracker Red CMX-ROS was used to determine ΔΨ_m_ in control untreated cells (0 μM MT) and in cells treated with 1 μM of MitoTam for one hour (1 μM MT). Staining with DAPI was used as an internal control for the analysis. Mean signal intensities of Mitotracker Red CMX-ROS and DAPI are depicted on the y-axis (mean ± s.d., ****p<0.0001). **I)** *In situ* ΔΨ_m_ was measured in digitonin-permeabilized BSF *T. brucei* cells stained with Safranin O dye. Where indicated, the F_o_F_1_-ATP synthase substrate – ATP, the F_o_F_1_-ATP synthase inhibitor - oligomycin (Olgm, green line) and MitoTam (1 µM, red line) were added. **J)** Red mitochondrial superoxide indicator MitoSOX was used to detect intramitochondrial ROS production in untreated BSF *T. brucei* control cells (-MT, blue) and BSF cells treated with 100 nM MitoTam for 16 hours (+MT 16 h, red) and 24 hours. Data were normalized to the values of the untreated control cells and expressed as a percentage (+MT 24 h, red) (mean ± s.d., *n* = 6, ****p <0.0001). **K)** Integrity of the inner BSF *T. brucei* mitochondrial membrane was assessed in isolated mitochondria incubated with increasing concentration of MitoTam. The mitochondrial enzyme threonine dehydrogenase (TDH) was used as a marker for permeability of the membrane, as the TDH activity is detectable only if a substrate passes freely through compromised mitochondrial membrane (mean ± s.d., *n* = 3).

To examine whether MitoTam affects mitochondrial activity, we analyzed changes in several cellular attributes associated with mitochondrial function. *T. brucei* BSF cultures were treated for 16 and/or 24 h with 40 nM or 100 nM of MitoTam. We found that incubation with 100 nM MitoTam significantly reduced glycerol-3-phosphate-dependent respiration, increased cellular ADP/ATP ratio (by 2.6-fold), and decreased mitochondrial and cytosolic ATP as well as total cellular ATP levels (Fig. 4B-E). Next, we assessed ΔΨ_m_ in live cells using the cell-permeable red fluorescent dye TMRE and using the Safranin O assay in digitonin-permeabilized cells. The results of both assays demonstrate that treatment with 100 nM MitoTam rendered the cell incapable of maintaining ΔΨ_m_ (Fig. 4F,G). Moreover, using mitochondrial superoxide indicator (MitoSOX), we detected a significant decrease of mitochondrial ROS levels upon MitoTam treatment, suggesting that the mode of action of MitoTam in trypanosomes differs from its effect in cancer cells (Fig. 4J). Experiments with cultures treated with MitoTam at 40 nM showed a similar but less pronounced overall effect on *T. brucei* mitochondrial activity (Fig. S3).

In agreement with these results, fluorescent microscopy evaluation revealed that treatment with 1 μM MitoTam for 1 h led to an ⁓8-fold decrease of the signal intensity of MitoTracker Red CMX-ROS, a mitochondrial probe used to stain live cells that relies on ΔΨ_m_ (Fig. 4H).

Furthermore, ΔΨ_m_ was assessed in digitonin-permeabilized wild-type BSF cells by Safranine O dye upon addition of ATP. We found that ΔΨ_m_ was quickly dissipated with 1 µM MitoTam, a phenotype dentical to the effect of addition of oligomycin, an inhibitor of ATP synthase (Fig. 4I). Moreover, using mitochondrial superoxide indicator (MitoSOX), we detected a significant decrease of mitochondrial ROS levels upon MitoTam treatment, suggesting that the mode of action of MitoTam in trypanosomes differs from its effect in cancer cells (Fig. 4J).

While the observed changes in the evaluated attributes are consistent with inhibition of FoF1-ATP synthase that generates ΔΨ_m_ in BSF *T. brucei* cells [46], we also tested if the rapid drop in ΔΨ_m_ is caused by non-specific disruption of the IMM. To this end, we assessed the integrity of isolated mitochondria using the mitochondrial marker threonine dehydrogenase (TDH), whose activity is detectable only if the added substrate threonine can freely pass through ‘compromised’ mitochondrial membrane. As shown in Fig. 4K, treatment with MitoTam at 7.4 µM decreased mitochondrial membrane integrity by 50%. This relatively high value (369 times higher than EC_50_) determined in this *in vitro* experiment can be explained by a need to form much larger membrane pores for threonine (molecular weight 119.12 g/mol) to enter mitochondria when compared to the formation of much smaller or temporal pores allowing the passage of H^+^. In addition, in this experiment, the BSF form mitochondria is not energized due to the lack of substrates and therefore this could have prevented accumulation of MitoTam from in the organelle. Taken together, these data indicate that MitoTam rapidly disrupts the integrity of the IMM, which leads to the ΔΨ_m_ collapse and cell death.

## Discussion

In cancer cells, MitoTam was shown to inhibit CI-dependent respiration, which leads to a rapid decrease of ΔΨ_m_, increased ROS production, and disruption of respiratory supercomplexes. These effects ultimately trigger cell death. The phenotype is even more pronounced in cancer cells, which exhibit high levels of the Her2 protein in mitochondria and elevated CI-dependent respiration. Consistently, mammalian cells deficient in CI are more resistant to MitoTam [19]. In metabolically active senescent cells, MitoTam treatment led to increased levels of ROS, reduced ΔΨm, impaired mitochondrial morphology, and metabolic switching to glycolysis for cellular ATP generation [22]. Interestingly, the mitochondrial phenotype was alleviated and cell survival prolonged in MitoTam- treated senescent cells overexpressing adenine nucleotide translocase-2 (ANT2), which imports ATP into mitochondria [22]. Furthermore, restriction of glycolytic pathways, either by limiting substrate levels or by adding glycolytic inhibitors competing with glucose sensitized control cells to MitoTam [22]. In contrast to cancer cells, ROS scavengers failed to rescue MitoTam-treated senescent cells from death. Moreover, treatment with established CI inhibitors (rotenone and piericidin A) alone did not selectively eliminate senescent cells, while they are sensitive to MitoTam treatment [22]. Therefore, the senolytic effect of MitoTam appears to involve additional mechanisms to CI inhibition. Taken together, the effect of MitoTam on cancer and senescent cells seems to be complex and includes targeting of CI, mitochondrial membrane depolarization, loss of ATP, increased levels of ROS, and possibly interplay between ATP/ADP exchange mechanism and ATP synthase [19, 22].

In this study, we showed that MitoTam inhibits the growth of a variety of parasitic protists and fungi. Different sensitivity to the compound may reflect different metabolic adaptations that the pathogens have developed to thrive in their host organism. High potencies were determined for *T. brucei* BSF and for the erythrocytic stage of *P. falciparum*, the relevant life cycle stages persisting in the mammalian host of both these medically important parasites. Notably, canonical CI is missing in *P. falciparum* [49] and is dispensable in BSF *T. brucei* [48, 50] suggesting that MitoTam has additional molecular targets besides CI with its NADH dehydrogenase. The EC_50_ value of MitoTam was one or two orders of magnitude higher (but still in in the micromolar range) for other tested organisms such as *Leishmania*, *Acanthamoeba* spp. and the widespread pathogenic yeasts *C. albicans* and *C. neoformans*. MitoTam was the least active against anaerobic protists *Trichomonas vaginalis* and *Giardia intestinalis*, likely due to the absence of conventional energized mitochondria in these organisms [33]. The low activity observed against free-living parasitic amoebae *Naegleria fowleri* requires further research.

Following the encouraging results from *in vitro* testing, we demonstrated that MitoTam significantly decreases parasitemia levels, as well as frequency and size of lesions in animals infected with *T. brucei* and *L. mexicana*, respectively. In agreement with published data [19, 22] [23], repeated injection of MitoTam had no apparent adverse effect on BALB/c mice used in our studies.

To investigate the mode of action of MitoTam in unicellular pathogens, we performed comparative proteomic analysis using two distinct model organisms, *T. brucei* (BSF) and *C. albicans*. Only minor changes were detected in the *C. albicans* proteome upon MitoTam treatment, the most upregulated proteins being two homologues of multidrug efflux pumps, possibly explaining the 28 times higher EC_50_ value when compared to BSF *T. brucei* cells. The corresponding analysis in *T. brucei* cells revealed substantial proteome changes upon 12 h MitoTam treatment. Whilst the complexity of the observed alterations failed to pinpoint the exact molecular target(s) of MitoTam, there was a significant impact on the mitochondrion.

To get insight into the complex mode of action that is linked to the function of CI in cancer cells [19], we decided to examine the effect of MitoTam on mitochondrial parameters in BSF *T. brucei*, where the respiratory complex is dispensable [48]. Consistent with studies on cancer and senescent cells [19, 22] [23], we found that cells incubated with MitoTam exhibit increased levels of cellular ROS and resulted in rapid dissipation of ΔΨ_m_, which leads to cell death. In contrast, mitochondrial superoxide levels in *T. brucei* cells were decreased upon treatment with MitoTam, indicating that the oxidative burst caused by inhibition of CI-dependent respiration is not responsible for *T. brucei* growth inhibition. Furthermore, *T. brucei* BSF cells incubated with MitoTam exhibited reduced glycerol-3- phosphate-stimulated oxygen consumption, decreased ATP levels, and increased ADP/ATP ratio, suggesting that the biochemical effect of MitoTam could either be due to inhibition of F_o_F_1_-ATP synthase that generates ΔΨ_m_ in these cells [45, 46] or, simply, due to disruption of the mitochondrial membrane integrity, resulting in ΔΨ_m_ collapse.

Our analysis shows that MitoTam at micromolar concentrations significantly increase the membrane permeability of isolated mitochondria, which could be the cause of the altered mitochondrial phenotype. This effect could be due to a direct interaction of the lipophilic MitoTam molecule with the membrane, causing unregulated proton leak, or perhaps due to alteration in membrane fluidity. Disruption of the membrane would indeed rapidly dissipate ΔΨ_m_ in live cells. Alternatively, the effect of MitoTam on membranes may be due to affecting enzymes involved in phospholipid synthesis. Intriguingly, studies on *T. cruzi*, *Leishmania spp*. and *P. falciparum* indicate that tamoxifen, the functional pharmacophore of MitoTam, has an inhibitory effect on phospholipid metabolic pathways [51–53]. Additional experiments are needed to reveal the multifactorial MitoTam mode of action. Nevertheless, our study shows that MitoTam represents a promising candidate to be repurposed as an effective anti-parasitic and anti-fungal compound with high selectivity.

## Material and Methods

### Pathogen culture

Culture conditions for all organisms used in this study are summarized in Supplementary Table 4

### Drug sensitivity assays

The cytotoxicity effect of MitoTam was tested in a variety of organisms. Briefly, cells were seeded in 96-well plates at concentrations summarized in Table 4 and grown under standard conditions for 24- 120 h, as indicated. The drug was serially diluted in a medium using a two-fold dilution across a plate, resulting in a total volume of 200 µl per well. Each experiment included untreated control cells, as well as a positive control treated with a known growth inhibitory compound for each respective pathogen. The results were statistically analyzed, and dose-response curves plotted using Prism 6.01 (GraphPad Software) and expressed as the half-maximal effective concentration (EC_50_).

The growth of *C. albicans* and *C. neoformans* cultures was determined from OD_600_ values measured on the I-Mark microplate reader (BioRad). *Plasmodium falciparum* dose-response curves were generated using SYBR Green I, as described previously [54]. In brief, triplicate two-fold compound dilution series were set up in 100 μl parasite cultures (̴0.2% parasitemia in 2% hematocrit). After two cycles (96 h), parasites were lysed with 20 μl of 6× SYBR green I lysis buffer (0.16% saponin; 20 mM Tris-HCl, 5 mM EDTA, 1.6% Triton X-100, pH 7.4), supplemented with 1:1000 SYBR green I (from 10000× stock, Thermo Fisher). Fluorescence intensity was measured with an Infinite M200 Pro Multimode Microplate Reader (Tecan). *Trypanosoma cruzi* epimastigote cell growth was assessed by CellTiter-Glo 2.0 cell viability assay in a 96-well plate according to the manufacturer’s protocol. As indicated, the dose-response curves of *T. brucei* and other pathogens were plotted from cell counts performed on the Guava EasyCyte 8HT flow cytometer (Luminex). The instrument setting and gate selection were adjusted using untreated control for each organism. For *G. intestinalis, A. castellanii,* and *N. fowleri,* that tend to adhere to the walls of the cultivation vessel, were placed on ice for 20 min and subsequently paraformaldehyde was added to a final concentration of 2% before counting the cells on the flow cytometer.

### Trypanosoma brucei growth analysis

*T. brucei* bloodstream cells were seeded at a concentration of 2×10^5^ and 8×10^5^ cells/ml of 5 ml HMI-9 medium in aerobic culture flasks. Biological triplicates were analyzed including untreated control cultures and cultures treated with final concentrations of 40 nM and 100 nM MitoTam, respectively. Cells were grown under standard growth conditions (37°C, 5% CO_2_). At each time point (8, 12, 16, 24, and 48 h), 20 µl were sampled, diluted 10x in fresh HMI-9 and the culture concentration was assessed using Guava EasyCyte 8HT flow cytometer (Luminex). The count parameters and gates were set according to previously measured cultures. In parallel, cells were checked under the microscope at each time point to confirm that live, moving cells were present in each culture.

### Trypanosoma brucei dead cell staining

*T. brucei* BSF cells were seeded at a concentration of 2×10^5^ cells/ml in 5 ml HMI-9 medium in aerobic cultivation flasks. Biological triplicates were set up, including an untreated control group and groups treated with final concentrations of 40 nM and 100 nM MitoTam. The cells were grown under regular growth conditions. At each time point (8, 12, 16, and 24 h), 20 µl were sampled from each flask was sampled, 10x diluted in fresh HMI-9 to a final volume of 200 µl and 2µl of SYTOX Green Dead Cell Stain (Invitrogen) was added to each sample. Cells were incubated at standard growth conditions for 30 min, after which live cell counts were analyzed using Guava EasyCyte 8HT flow cytometer (Luminex). The count parameters and gates were set according to previously measured cultures.

### Measurement of ROS production in *T. brucei*

Intracellular and intramitochondrial ROS production was monitored using published protocols [55]. Intracellular ROS levels were evaluated using three biological replicates of approximately 1×10^6^ untreated cells and cells treated with 100 nM MitoTam for 12 h. The samples were collected, incubated with 10 µM of 2′,7′-dichlorofluoresceine diacetate (DCFH-DA, Sigma-Aldrich) for 30 minutes and washed with PBS-G (1x PBS with 6 mM glucose). Using a 488 nm excitation laser and a 530/30 nm detector, 10,000 events were immediately counted on the BD FACSCanto II instrument.

Mitochondrial ROS production was assessed using the MitoSOX indicator (Thermo Fisher Scientific) in untreated cells and cells treated with 40 nM or 100 nM MitoTam. An equal number of cells (1×10^6^) for each treatment was collected, washed in PBS-G, resuspended in HMI-9 medium with 5 μM MitoSOX and stained for 30 min at 37°C, 5% CO_2_. After staining, cells were spun down, resuspended in PBS, and immediately analyzed by flow cytometry (BD FACS Canto II Instrument, 488 nm excitation laser and 585/15 nm emission). For each sample, 10 000 events were collected. All values were plotted and statistically analyzed using Prism (8.0) (GraphPad Software).

### Comparative label-free proteomics

For *T. brucei* bloodstream forms, a final concentration of 100 nM MitoTam was added to approximately 3×10^7^ cells in the exponential growth phase and cultures incubated for 14 h. Cell viability was checked microscopically before harvesting and washing three times in PBS (1200g, 10 min, 4°C). An untreated control group was prepared in parallel. Both groups were prepared in three biological replicates. The pellets were subjected to reductive alkylation and tryptic digest using routine procedures. Peptides were then analysed by liquid chromatography-tandem mass spectrometry on an ultimate3000 nano rapid separation LC system (Dionex) coupled to an Orbitrap Fusion mass spectrometer (Thermo-Fisher Scientific). Data were analysed as described in [43, 56] using label-free quantification in MaxQuant [41] searching the TriTrypDB *T. brucei* strain TREU927 protein database

For *C. albicans*, approximately 1×10^4^ cells were inoculated in RPMI media, as described in [58] and left to grow for 24 h at 35°C without agitation. MitoTam was added to the final concentration of 4.4 µM and cultures incubated for 12 h. Cells were then checked under a microscope, harvested, washed three times with PBS (1 200g, 10 min, 4°C). Pellets were then lyzed using FastPrep 24 5G homogenizer (MP Biomedicals) according to the manufacturer’s instructions. Lysates were used for the label-free proteomic analysis and compared with the untreated control group prepared in parallel. Both groups were prepared in three biological replicates. Analysis was performed based on the *C. albicans* protein database downloaded from Uniprot on 12.11.2019 [59]. The threshold settings for comparative proteomic analyses and data processing were chosen as described in [43] for both organisms. Briefly, thresholds were set to Q-value = 0, unique peptides detected >2 and the protein had to be identified at least twice in one of the conditions. For proteins found under only one condition, the average log2 converted intensity of 23 was selected as a minimum threshold. Significantly changed proteins were considered only if the fold change was >2.0.

In the proteomic results of *C. albicans,* localization was predicted based on Uniprot [59]. For *T. brucei* proteomic analysis, manual annotation and localization prediction were based on *T. brucei* 927 database or *T. brucei* bloodstream form mitoproteome database obtained from [44]. Volcano plots were drawn in Perseus 1.6.15.0 [42]. At first, proteins only identified by site, reverse, potential contaminants, and proteins detected in less than two experiments within at least one group were excluded. The imputation was performed using a normal distribution with width 0.3 and downshift of 1.8, separately for each column. The parameters of the volcano plot were set up as following: statistical t-test for both sides, 250 randomizations, false discovery rate of 0.05 and S_0_ of 0.1.

### Mitochondrial membrane potential measurements

The mitochondrial membrane potential (ΔΨm) of BSF *T. brucei* cells was estimated using the red fluorescent stain MitoTracker Red CMXRos (Invitrogen). MitoTam was added to 5 ml *T. brucei* culture in an exponential growth phase at a final concentration of 1 μM and cells were incubated for one hour under standard conditions. Subsequently, MitoTracker Red CMXRos was added to a final concentration of 100 nM and samples were incubated for another 10 min. Cultures were spun (1,300 g, 10 min, RT), resuspended in 5 ml of growth medium with 1 µM MitoTam and left for an additional 20 min under standard growth conditions. An untreated control was prepared in parallel. Treated and untreated cells were washed and transferred to 300 µl PBS, spread on microscopy slides, and left to settle. The slides were fixed by immersion in ice-cold methanol for 10 min, once the excess methanol had evaporated, the slides were mounted using Vectashield with DAPI (Vector laboratories), covered with cover slides, and sealed using nail varnish. The slides were imaged on a Leica TCS SP8 WLL SMD-FLIM microscope, using LAS X 3.5.6 imaging software. All images were captured using the exact same microscope setting. Quantification and comparison of signals were performed in NIS- Elements 5.30 (Nikon). Briefly, 10-20 z-stacks were captured using LAS X automatic settings. A single stack with the highest overall intensity was chosen from the batch, and using the Artificial Intelligence module, areas of signal corresponding to individual cells were mapped and the average intensity was determined from the selected area. Furthermore, cell DNA visualized by DAPI was also quantified in the same manner and compared across all images as a reference value. The signal intensity of MitoTracker Red CMXRos was then compared in treated and untreated control, while the intensity of the DAPI signal was used to confirm the reproducibility of the reading.

Estimation of ΔΨm in live cells was performed as described previously [55]. Briefly, an equal number of cells (1×10^6^) for each treatment was spun (1,400 g, 10 min, RT), the pellets were resuspended in HMI-9 medium with 60 nM TMRE (Thermo Fisher Scientific T669) and stained for 30 min at 37°C, 5% CO_2_. After staining, cells were spun down (1,400 g, 10 min, RT) and resuspended in 2 ml of PBS and immediately analyzed by flow cytometry (BD FACS Canto II Instrument). For each sample, 10,000 events were collected. As a control for mitochondrial membrane depolarization, cells were treated with 20 μM protonophore FCCP (carbonyl cyanide 4-(trifluoromethoxy) phenylhydrazone, Sigma). Data were evaluated using BD FACS Diva software (BD Company) and further analyzed using Prism (8.0) (GraphPad Software).

Estimation of the ΔΨm was also performed using the fluorescent probe Safranine O (Sigma) according to [55]. From each treatment, 2×10^7^ cells were collected, spun down (1,400 g, 10 min, RT), and resuspended in ANT buffer (8 mM KCl, 110 mM potassium gluconate, 10 mM NaCl, 10 mM HEPES, 10 mM K_2_HPO_4_, 0.015 mM EGTA, 0.5 mg/ml fatty acid-free BSA, 10 mM mannitol, 1.5 mM MgCl_2_) with 5 μM Safranine O. Intact live cells were permeabilized by the addition of 4 µM digitonin (Calbiochem). The fluorescence of all samples was measured at RT and constant stirring and recorded on a Hitachi F-7100 spectrofluorometer (Hitachi High Technologies) at a 5 Hz acquisition rate, using 495 and 585 nm excitation and emission wavelengths, respectively. As indicated, the reaction was started by adding 1 mM ATP (PanReac AppliChem), F_o_F_1_-ATP synthase substrate, and stopped by the addition of 10 µg/ml oligomycin (Sigma), F_o_F_1_-ATP synthase inhibitor. The protonophore SF6847 (Enzo Life Sciences) was added at a final concentration of 250 nM as control for maximal depolarization. Fluorescence data were analyzed using Prism (8.0) (GraphPad Software).

### High-resolution respirometry

The effect of MitoTam on respiration was analyzed by Oroboros Oxygraph-2K (Oroboros Instruments Corp., Innsbruck, Austria) as described [55]. Bloodstream form *T. brucei* cells were incubated with 40 nM or 100 nM MitoTam for 16 h and 24 h, as indicated. For each treated sample and control, 2×10^7^ cells were spun down (1,400 g, 10 min, RT) and pellets were washed in Mir05 medium (0.5 mM EGTA, 3 mM MgCl_2_, 60 mM lactobionic acid, 20 mM taurine, 10 mM KH_2_PO_4_, 20 mM HEPES, 110 mM sucrose, 1 mg/ml fatty acid-free BSA, pH 7.1). Before the measurement started, the pellets were resuspended in 0.5 ml of Mir05 medium preheated to 37°C and transferred to the respiration chamber. Respiration was monitored at 37°C and with constant stirring. The experiment started with the addition of 10 mM glycerol-3-phosphate (Sigma), the mitochondrial glycerol-3-phosphate dehydrogenase substrate, and respiration was inhibited by the addition of 250 μM SHAM (Salicylhydroxamic acid), the inhibitor of the trypanosomal alternative oxidase. The acquired data were analyzed using Prism (8.0) (GraphPad Software).

### Western blot analysis

*T. brucei* BSF cells were incubated with 100 nM MitoTam and harvested at different time points after addition as indicated (0h, 8h, 12h, 16h and 24h). Cells were spun down (1,400 g, 10 min, RT), pellets were washed in PBS (pH 7.4) and resuspended in 3xSDS-Page sample buffer (150 mM Tris pH 6.8, 300 mM 1,4-dithiothreitol, 6% (w/v) SDS, 30% (w/v) glycerol, 0.02% (w/v) bromophenol blue). The whole-cell lysates were denatured at 97°C for 8 min and stored at -80°C. For Western blot analysis, a volume of sample equal to 2.5×10^6^ cells per well was loaded onto 12% gel, separated by SDS-Page, blotted onto a nitrocellulose membrane (Amersham Protram 0,2 µm PC GE Healthcare Life Sciences) and probed with a monoclonal (mAb) or polyclonal antibody (pAb). Incubation with primary antibodies was followed by a secondary antibody, either HRP-conjugated goat anti-rabbit or an anti-mouse antibody (1:5,000, BioRad). Antibodies were detected using the enhanced chemiluminescence system (Immobilon Forte Western HRP substrate, Merck) on the Amersham Imager 600 (GE Health Care Life Sciences). The PageRuler™ Plus prestained protein ladder (Thermo Fisher Scientific 26619) was used to determine the size of the detected bands. Primary antibodies were: mAb anti-AOX (1:1,000), pAb anti-AAC (1:1,000), anti-VDAC (1:1,000), pAb anti-PYK (pyruvate kinase) 1:1,000 and antibodies against F_o_F_1_-ATP synthase subunits β (1:2,000), p18 (1:1,000), Tb1 (1:1,000), Tb2 (1:1,000), and OSCP (1:1,000) [55, 60].

### Measurement of the ATP/ADP ratio and total cellular ATP levels

Changes in the ATP/ADP ratio and total cellular ATP were determined in BSF *T. brucei* cells using the D-luciferin-luciferase bioluminescent enzymatic reaction (assay kit Sigma MAK135) according to manufacturer’s instructions. Briefly, from each sample 1×10^6^ cells were harvested and washed once with PBS supplemented with 6 mM glucose (PBS-G). Pellets were resuspended in PBS-G and the mixture transferred into a microtiter plate (96-well white flat-bottom). The bioluminescence signal was recorded in an Orion II microplate luminometer (Titertek Berthold) and analyzed using Prism (8.0) (GraphPad Software).

### *In situ* measurement of ATP levels

Cytosolic and mitochondrial ATP was measured using BSF *T. brucei* cell lines expressing firefly luciferase with or without N-terminal mitochondrial localization signal (MLS) following published protocols [61]. Briefly, MitoTam treated cells, as well as untreated control, were spun (1×10^7^ cells) (1,400 g, 10 min, RT) and washed in PBS-G. The pellets were resuspended in HEPES-LUC+Glu buffer (10 mM D-glucose, 20 mM HEPES, 116 mM NaCl, 5.6 mM KCL, 8 mM MgSO_4_, 1.8 mM CaCl_2,_ pH 7.4) and transferred to a 96-well microtiter plate. ATP-dependent luciferase bioluminescence was recorded on a plate reader (Tecan Infinite M100). The light emission was started with the addition of a D-luciferin solution (50 μM; Sigma), collected for 5 min, and statistically analyzed using Prism (8.0) (GraphPad Software).

### Determination of the mitochondrial membrane integrity

*T. brucei* bloodstream mitochondria were isolated by digitonin fractionation, according to [62]. Briefly, approximately 1×10^8^ cells were harvested (1400 g, 10 min, RT) and transferred to Hank’s balanced salt solution (Sigma Aldrich). The protein concentrations of the samples were determined by the BCA assay kit (Sigma-Aldrich, USA). Digitonin (Calbiochem) was added to a protein:mass ratio of 1:0.15, lysate was incubated for 4 min and then spun. The pellet was washed and used as a mitochondria-enriched fraction, while the supernatant was used as a cytosolic fraction. Activities of two marker enzymes were used to assess the purity of fractions and intactness of mitochondrial membranes. The enzyme activities of cytosolic pyruvate kinase (PYK) and mitochondrial threonine dehydrogenase (TDH) were assayed spectrophotometrically at 340 nM by monitoring NADH concentration changes during the reaction. The activity of PYK was measured according to [63] in 0.1M TEA buffer (ThermoFisher Scientific) (final pH 7.6), 5 mM MgSO_4_ and 50 mM KCl, with the addition of 2.8 mM phosphoenolpyruvate, 2 mM ADP, 0.3 mM NADH and lactate dehydrogenase. The activity of TDH was measured in 0.2M Tris-HCl buffer with 0.25 M KCl (final pH 8.6), with the addition of 120 mM threonine and 2.5 mM NAD^+^. Different concentrations of MitoTam were added to the reaction and TDH activity was monitored as a marker for disruption of the mitochondrial membranes. Treatment with nonionic detergent Triton-×100 was used to completely disrupt mitochondrial membranes and thus release maximum TDH activity. Data were statistically analyzed and plotted with Prism (8.0) (GraphPad Software).

### Survival analysis in a mouse model

To evaluate the trypanosomiasis effect of MitoTam on the mortality in mice, a group of 14 BALB/c mice was infected IP with approximately 2×10^5^ BSF *T. brucei* cells strain STIB920 and the group of 14 mice was infected with 5×10^5^ BSF cells of *T. brucei brucei* 427 strain, both in 100 µl sterile PBS. Mice were monitored every 12 hours. Based on previous experience with the progression of the disease, half of the infected mice from each group were IV injected with MitoTam (3 mg/kg bw) on days 3 and 5 after infection. The survival of the mice was visually monitored for up to 13 days after infection, the day of death was recorded for each animal.

### *T. brucei* mouse infection model

To assess the effect of MitoTam on the morbidity of trypanosomiasis in mice, sixteen BALB/c mice were infected with 2×10^5^ BSF *T. brucei* cells (strain STIB920) in 100 µl of sterile PBS. On day 2 and 4 post infection, eight mice were injected with MitoTam at a final concentration of (3 mg/kg bw).

Mice were monitored daily, and blood samples were collected from a tail prick from each animal. Blood drops were smeared on microscopy slides and stained using Wright-Giemsa stain modification, Diff-Quick (Medion Diagnostics) according to the manufacturer’s protocol. From each slide three images of random places, where red blood cells were not overlaying, were taken using an inverted widefield microscope Eclipse Ti2 (Nikon) using the CFI Plan Apochromat Lambda 20x objective (Nikon) with NIS-Elements AR 5.20 (Nikon). The images were then processed using the Artificial Intelligence module on NIS-Elements 5.30 (Nikon), manually trained to detect and quantify the number of red blood cells and trypanosomes. The ratio of trypanosomes to red blood cells was calculated and used to plot the data and calculate the statistical difference (student t-test) between the treated and untreated groups in a given days. Data were plotted with Prism (8.0) (GraphPad Software). Animal handling was approved by the Czech Ministry of Agriculture (53659/2019-MZE-18134).

### *L. mexicana* mouse subcutaneous leishmaniasis model

The culture conditions of the *L. mexicana* promastigotes were as indicated in Supplementary Table 4, their concentration was determined by hemocytometer. Before infection experiments, promastigotes were harvested, washed three times and resuspended in PBS. Twenty BALB/c mice (ten weeks old) were anesthetized IP with a mixture of ketamine (150 mg/kg) and xylazine (15 mg/kg) and infected intradermally in the ear pinnae by injection of 10^7^ promastigotes. For a group of ten animals, MitoTam was administered IP at 1, 2, 3, 5, 6 and 7 weeks after infection at a dose of (3 mg/kg body weight), other ten animals served as nontreated controls. The presence of lesions was monitored for 12 consecutive weeks.

### Rescue assay for *Leishmania spp.* amastigotes

Initially, murine macrophage cells (J774) were seeded in RPMI supplemented with 10% FCS and 50 µg/ml phorbol 12-myristate 13-acetate and left to differentiate for 24 h. Then, after washing the cells once with warm (37°C) serum-free RPMI, stationary phage promastigotes of *L. infantum* or *L. major* were added in a 10:1 ratio (2.5×10^6^ cell/ml). After 24 h of incubation, cells were washed five times with serum-free RPMI and exposed to increasing concentration of MitoTam (or amphotericin B, as positive control) in RPMI (2% FCS). After two days, macrophages were lysed with 20 µl of 0.05% SDS in RPMI for 30 s and released *Leishmania* cells were incubated with M199 10% FCS for another three days. Finally, the viability of transformed live cells was determined by the resazurin viability assay.

## Acknowledgment

The project was supported by the Czech Science Foundation (20-28072S) to RS and 20-14409S to AZ, CePaViP provided by The European Regional Development Fund and Ministry of Education, Youth and Sports of the Czech Republic (reg. no. CZ.02.1.01/0.0/0.0/16_019/0000759) to RS, PV, MZ and AZ, Grant Agency of Charles University (406722) to DA and Fundação para a Ciência e Tecnologia (FCT, Portugal)—PD/BD/128002/2016 for providing funds to MM. We thank to Karel Harant and Pavel Talacko from the Laboratory of Mass Spectrometry, Biocev, Charles University, Faculty of Science, where proteomic and mass spectrometric analysis had been done.

## Supplementary data

**Table S1: Table of screened pathogens summarizing their impact on global health.**

**Table S2: *C. albicans* proteomic analysis after 12 h of MitoTam exposure.**

Proteome comparison of *C. albicans* cultivated in the presence of 4.4 µM MitoTam for 12 h and untreated culture. The table is organized in four sheets: All detected proteins, Downregulated in MitoTam, Upregulated in MitoTam and Raw data. The first three sheets are showing fold-abundance change only for clarity. Upregulated and downregulated proteins were filtered by >2-fold change. Proteins were identified and, where applicable, subcellular localization was annotated based on Uniprot [59].

**Table S3: *T. brucei* proteomic analysis after 14h of MitoTam exposure.**

Proteome comparison of *T. brucei* cultivated in the presence of 100 nM MitoTam for 14 h and untreated culture. Table is organized in four sheets: All detected proteins, Downregulated in MitoTam, Upregulated in MitoTam and Raw data. The first three sheets are simplified to demonstrate fold-abundance change only for clarity. Upregulated and downregulated proteins were filtered by >2-fold change. Where applicable, manual annotation and localization prediction was based on the *T. brucei* 927 mitochondrial proteome [44].

**Table S4: Summary of culture conditions and viability assays for all organisms used in this study.**

**Figure S1: *Leishmania* sensitivity towards MitoTam evaluated by the intramacrophage assay.**

Murine macrophage cell culture J774A.1 was infected with amastigotes of *L. major* (pink line) or *L. infantum* (brown line) and incubated with increasing concentration of MitoTam as indicated on the x-axis. Uninfected macrophages (green line) were included as a control.

**Figure S2: Survival analysis of BALB/c mice infected with *T. brucei*.**

Survival of infected mice was monitored daily for 8 days. As depicted (dashed lines) half of the infected mice were injected with MitoTam (+MitoTam) on days 2 and 4 post-infection (3 mg/kg body weight). Number of surviving mice from MitoTam treated group (red line) as well as from the untreated control group (-MitoTam) (blue line) are depicted.

**Figure S3: MitoTam alters the mitochondrial function of bloodstream *T. brucei*.**

Cells were incubated with 40 nM of MitoTam for 16 and 24 hours and their mitochondrial parameters were assayed as for 100 nM MitoTam.

**A)** The O_2_ flux per cell using high-resolution respirometry after addition of glycerol-3-phosphate was determined in BSF *T. brucei* untreated control cells (-MT, blue) and BSF cells treated with 40 nM MitoTam for 16 hours (+MT 16 h, red) and 24 hours (+MT 24 h, red) (mean ± s.d., *n* = 3, **p < 0.01, ***p < 0.001).

**B)** Relative ADP/ATP ratio analyzed using a bioluminescence assay kit in BSF *T. brucei* untreated control cells (-MT, blue) and BSF cells treated with 40 nM MitoTam for 16 hours (+MT 16 h, red) and 24 hours (+MT 24 h, red) (mean ± s.d., *n* = 3).

**C)** Cytosolic and mitochondrial ATP levels were assessed in transgenic BSF *T. brucei* cell lines expressing firefly luciferase. Results of untreated control cells (-MT, blue) were compared with results of cultures treated with 40 nM MitoTam for 16 hours (+MT 16 h, red) and 24 hours (+MT 24 h, red). Data were normalized to the respective values of the untreated control cells and expressed as a percentage (mean ± s.d., *n* = 4).

**D)** Total cellular ATP levels in BSF *T. brucei* were determined using a bioluminescence assay kit. Data from cultures treated with MitoTam for 16 hours (+MT 16 h, red) and 24 hours (+MT 24 h, red) were normalized to the values of the untreated control cells (-MT, blue) and expressed in percentage (mean ± s.d., n = 4).

**E)** Flow cytometry of TMRE stained cells was used to determine ΔΨ_m_ of BSF *T. brucei*. Data from cultures treated with 40 nM MitoTam for 16 hours (16 h +MT, red) and 24 hours (+MT 24 h, red) were normalized to the values of the untreated control cells (-MT, blue) and expressed in percentage. Uncoupler FCCP was added as a control for ΔΨ_m_ depolarization (mean ± s.d., n = 6, ****P <0.0001).

**F)** *In situ* ΔΨ_m_ was measured in digitonin-permeabilized BSF *T. brucei* cells stained with Safranin O dye. Where indicated, the F_o_F_1_-ATP synthase substrate – ATP, the F_o_F_1_-ATP synthase inhibitor - oligomycin (Olgm) and the protonophore SF6847 were added. Representative traces from measurement of untreated control cells (-MT, blue) in comparison with cells treated with 40 nM MitoTam for 16 hours (+MT 16 h, orange) and 24 hours (+MT 24 h, red) are shown.

